# First report of classical knockdown resistance (*kdr*) mutation, L1014F, in human head louse *Pediculus humanus capitis* (Phthiraptera: Anoplura)

**DOI:** 10.1101/2022.05.14.491638

**Authors:** Prashant K. Mallick, Ankita Sindhania, Toshi Gupta, Dhirendra P Singh, Seema Saini, Om P. Singh

## Abstract

There are at least three known knockdown resistance (*kdr*) mutations reported globally in the human head louse *Pediculus humanus capitis* (Phthiraptera: Anoplura) that are associated with reduced sensitivity to pyrethroids. However, prevalence of *kdr* mutation is not known in the Indian subcontinent. To identify *kd*r mutations in the Indian population, the genomic region of the voltage-gated sodium channel gene encompassing IIS1-2 linker to IIS6 segments was PCR-amplified and sequenced from *P. humanus capitis* samples collected from different geographic localities of India. DNA sequencing revealed the presence of four *kdr* mutations, i.e., M827I, T929I, L932F and L1014F. The presence of a classical *kd*r mutation, L1014F, a most widely reported mutation across insect-taxa associated with *kdr*-trait, is being reported for the first time in *P. humanus capitis*.

## Introduction

*Pediculus humanus capitis* (Phthiraptera: Anoplura), commonly known as ‘head louse’, is an obligate hematophagous ectoparasite of humans widely distributed globally. Pediculosis is very common, especially among children, even in developed countries. Head louse may cause substantial discomfort and irritability, social distress and embarrassment, sleeplessness and sores on the head caused by scratching which can sometimes become infected. Recent studies have reported molecular evidence of the presence of various pathogens in the *P. humanus capitis*; such as *Borrelia recurrentis*, the causative agent of relapsing fever (Amanzougaghene et al., 2016), *Bartonella quintana*, the causative agent of trench fever (Sasaki et al., 2006; Sunantaraporn et al., 2015) and Acinetobacter spp. (Louni et al., 2018). Head lice is also considered to be a potential vector of *Rickettsia prowazekii*, the aetiological agent of louse-borne epidemic typhus (Robinson et al., 2003).

The control of lice relies primarily on the use of over-the-counter products containing pyrethroids. Permethrin 1% or pyrethrins (pyrethroids) is recommended as a reasonable first choice for primary treatment of active infestations being relatively safe for mammals (Devore et al., 2015). Pyrethroids interact with the open state of the voltage-gated sodium channel (VGSC) in the nervous system of insects and cause a prolonged influx of sodium that leads to nerve depolarization and eventually death by muscle paralysis. Knockdown resistance (*kdr*) is one of the mechanisms of pyrethroid resistance in insects, which is conferred by one or more amino acid substitutions in the VGSC (target site) leading to reduced sensitivity to pyrethroids. Three *kdr* mutations, i.e., M827I, T929I and L932F (amino acid positions shown here are based on house-fly sequence), have been reported to be present in most parts of the world (Hodgdon et al., 2010; Mohammadi et al., 2021). Another mutation D11E was suggested to be not involved in insecticide resistance (Tomita et al., 2003). An exceptionally high number of novel non-synonymous mutations (seven mutations in head lice and 10 in the body lice) were recorded present in a small region of the VGSC (between IIS1-2 linker and IIS5) by a research group in Iran (Firooziyan et al., 2017; Ghahvechi Khaligh et al., 2021). However, there is no report of classic L1014F-*kdr* mutation so far in the head louse, which is the first mutation mapped to *kdr*-trait in the house fly *Musca domestica* (Williamson et al., 1996) and is the most commonly reported *kdr* mutation in a wide array of insects (Rinkevich et al., 2013; Dong et al., 2014). In some insects, this mutation is not found due to the codon constraint, where at least two nucleotide substitutions are required for a single amino acid substitution (Davies et al., 2007). In *Pediculus* sp., however, no such constraint exists associated with L1014. Because most of the earlier studies demonstrating the presence of *kd*r mutation in head lice populations, have focused on the already known three *kdr* mutations, and have ignored sequencing of L1014 residue, it is desirable to investigate the presence of this classical mutation in human louse. This study was therefore undertaken to investigate the prevalence of *kdr* mutations in Indian head louse populations with an emphasis on checking the probable existence of the classical mutation L1014F. To date, there is no study undertaken to detect *kdr* mutation in populations from countries in the Indian subcontinent.

## Methods

Head lice were collected from six different localities representing different parts of India, viz., from Village Marykulam (9° 69’ E, 77° 04’ N), district Idukki of Kerala state, representing southern India; Village Chandpur (19° 94’ E, 85° 41’ N), district Nayagarh of Odisha state, representing eastern India; Shashtri Nagar (26°96’ E, 75.79’ N), Jaipur city of Rajasthan state, representing western India; village Sapas (27°19’E, 81°51’ N) of district Bahraich, Jawahar Park (28°51’E, 77°23’N) colony of New Delhi, village Chadwal (32°47’ E, 75°32’ N), district Kathua of Jammu & Kashmir, representing northern India. The lice, which were dislodged from the hair of infested adult female volunteers (age 18-25 years) during routine combing, were stored in a microcentrifuge tube containing a piece of silica gel (provided by the study team) by themselves. The tube was collected by the study team and stored in a -20 °C deep freezer. No intervention of human subjects was involved. DNA was isolated from individual louse manually following the method described by Black WC (1997). For PCR amplification and sequencing of the IIS4-S6 segment of the VGSC, primers PdUF (5’-ACC CAT TCG TCG AAT TAT TCA TAA CT-3’) and PdUR (5’-ATG CTT CGT TTT ACC CAT GC-3’) were designed from exon 14 and 18, respectively. Six additional primers, i.e., PdE1F (5’-CAA CGT TTG CCA TTG AAG C-3’), PdE1R (5’-GTA ACA CGG AAA GCC CTT GA-3’), PdE2F (5’-AAA TCG TGG CCA ACG TTA AA), PdE2R (5’-CCA AAA AGT TGC ATT CCC ATA-3’), PdE3F (5’-GGA TAA AGA ACT TCC CAG ATG GA-3’), and PdE3R (5’-ACA ACA GTG GCC AGG AAA AA-3’) were designed for sequencing. The locations of primers have been shown in **Figure 1**. The PCR amplification was carried out on gDNA of lice in a reaction mixture of 20 µl volume with GoTaq Green Master Mix (Promega), 0.5 µM of each primers (PdUF and PdUR) and 1 µl of DNA template. The PCR conditions were: Initial denaturation at 95°C for 2 min, followed by 35 cycles each with 95°C for 30 sec, 55°C for 30 sec and 72°C for 1.5 min, and a final extension at 72°C for 7 min. Two µL of the PCR product were electrophoresed on 2% agarose gel, checked under UV-Gel Doc, and the remaining PCR products were purified with Exo-Sap IT (Thermo Fisher Scientific). Each PCR product was subjected to a sequence termination reaction by all eight primers. The products were capillary-electrophoresed in an in-house capillary DNA sequencer ABI Prism 3730xl. The basecalls in the sequence chromatogram were edited manually. Sequence reads which collapsed due to the presence of indels in intron regions, as evidenced by the presence of overlapping sequences (**Supplementary Figure S1**), were trimmed before performing alignment and further analysis.

**Figure 1.**
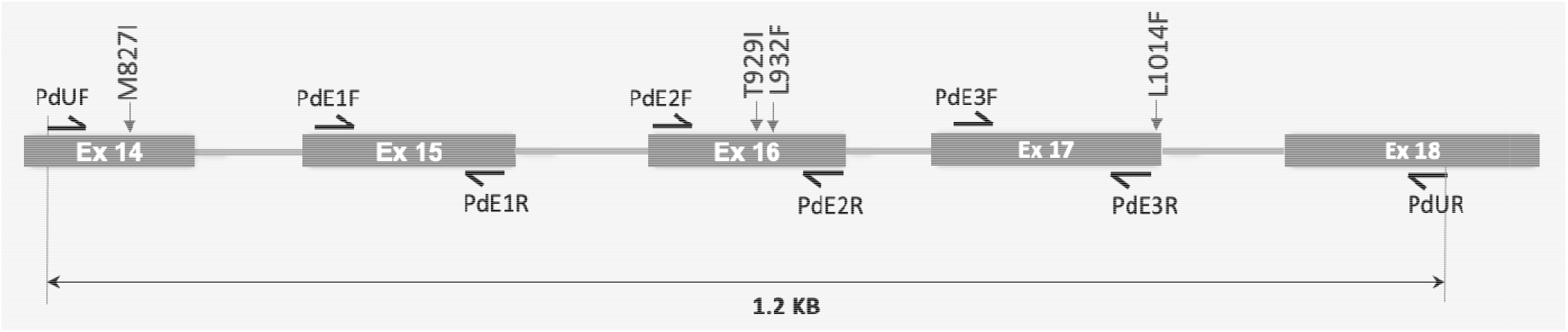
Schematic presentation of PCR amplification and DNA sequencing strategy. The flanking primers PdUF and PdUR were used for PCR amplification and all eight primers (denoted by harpoons) were used individually for sequencing termination reactions. The locations of *kdr* mutations have been indicated by vertical texts.

## Results

Four non-synonymous mutations, M827I (ATG→ATT), T929I (ACA→ATA), L932F (CTT→TTT) and L1014F (TTA→TTT) were recorded (**Figure 2**). The amino acid positions shown here are based on the house fly sequence that corresponds to M815I, T917I, L920F and L992F in *Pediculus* sp., respectively. The number of samples sequenced, genotyping results and allelic frequencies of *kdr* mutations in different populations have been shown in **Table 1**. Kerala population had the highest frequencies o all four *kdr* mutations with the presence of new mutation L1014F.

**Table 1.** Genotyping results of *kdr* alleles and their allelic frequencies in different populations

| Location (state) | N* | M827I |  |  | T929I |  |  | L932F |  |  | L1014F |  |  | Frequencies of mutant *kdr* alleles |  |  |  |
| --- | --- | --- | --- | --- | --- | --- | --- | --- | --- | --- | --- | --- | --- | --- | --- | --- | --- |
|  |  | MM | MI | II | TT | TI | II | LL | LF | FF | LL | LF | FF | 827I | 929I | 932F | 1014F |
| Kerala | 62 | 32 | 21 | 9 | 57 | 4 | 1 | 46 | 8 | 8 | 44 | 10 | 8 | 0.31 | 0.05 | 0.19 | 0.21 |
| Odisha | 19 | 17 | 1 | 1 | 19 | 0 | 0 | 17 | 0 | 2 | 19 | 0 | 0 | 0.08 | - | 0.11 | - |
| Rajasthan | 11 | 11 | 0 | 0 | 11 | 0 | 0 | 10 | 0 | 1 | 11 | 0 | 0 | - | - | 0.09 | - |
| UP* | 24 | 22 | 2 | 0 | 23 | 1 | 0 | 21 | 1 | 2 | 24 | 0 | 0 | 0.04 | 0.02 | 0.10 | - |
| Delhi | 3 | 1 | 1 | 1 | 3 | 0 | 0 | 3 | 0 | 0 | 3 | 0 | 0 | 0.50 | - | 0.00 | - |
| J&K* | 10 | 9 | 1 | 0 | 10 | 0 | 0 | 9 | 1 | 0 | 10 | 0 | 0 | 0.05 | - | 0.05 | - |
| Total | 129 | 92 | 26 | 11 | 123 | 5 | 1 | 106 | 10 | 13 | 111 | 10 | 8 | 0.19 | 0.03 | 0.14 | 0.10 |

\*Abbreviations used: N=number of samples; UP= Uttar Pradesh; J&K= Jammu & Kashmir

**Figure 2.**
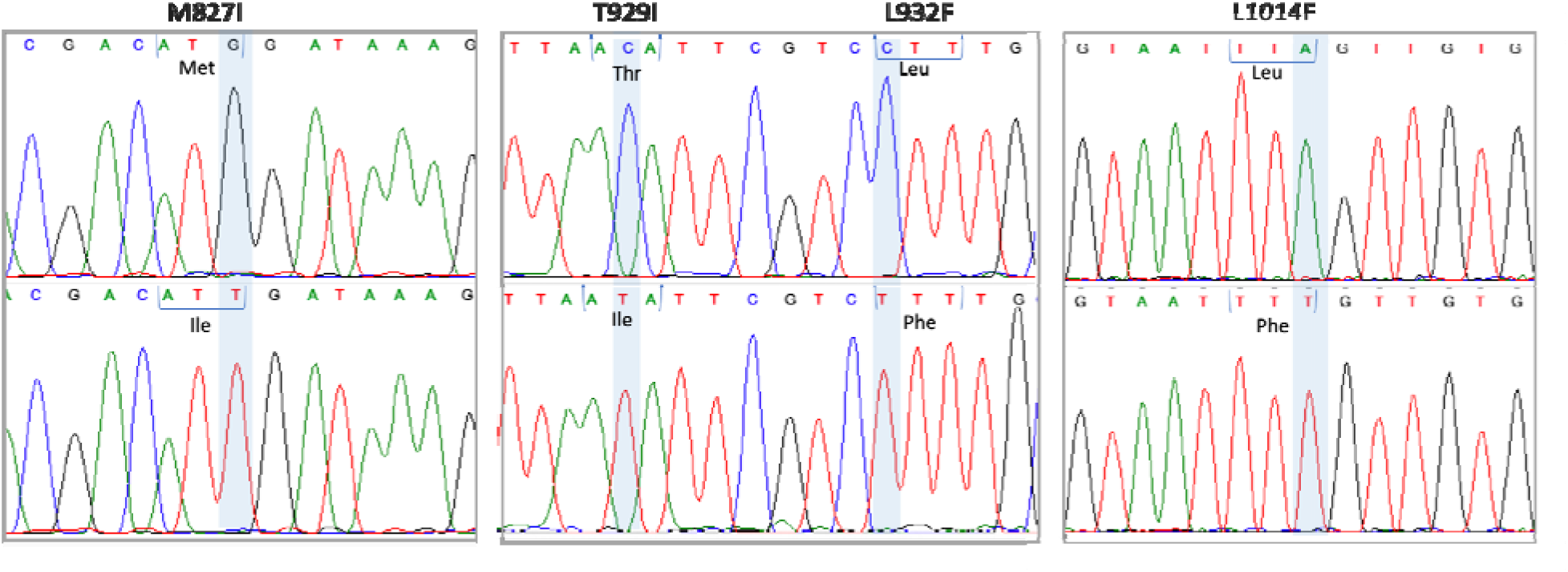
DNA sequencing electrophoretograms showing nucleotide substitutions and amino acid changes

## Discussion

Topical use of insecticides pyrethrins/pyrethroids-based formulations is the most common treatment for pediculosis, which dominates the over-the-counter market worldwide (Clark et al., 2013). Resistance to pyrethroids in head lice has been reported in several countries. Knockdown resistance is one of the mechanisms of pyrethroid resistance that renders pyrethroid-based pediculicides ineffective. Several *kdr*-mutations have been reported in the head louse (Hodgdon et al., 2010; Mohammadi et al., 2021), and the three most common mutations reported worldwide (M827I, T929I and L932F) have been shown to reduce sensitivity to permethrin, alone or together, when expressed in *Xenopus* oocyte (SupYoon et al., 2008). However, the T929I mutation, either alone or in combination, abolished permethrin sensitivity (SupYoon et al., 2008). In this study we recorded all of the above three mutations in the Indian population with varying frequencies along with a new mutation, L1014F, not reported earlier in head louse. The L1014F mutation was the first reported mutation in the VGSC associated with *kdr*-trait in *Musca domestica* and reported to be present in a wide array of insects. In some insects, e.g., *Aedes aegypti*, this mutation was never found due to constraints associated with the codon, which required a minimum of two SNPs to take place for a single amino acid substitution. In the head louse, however, there is no such constraint, which prompted us to screen this mutation along with the other three mutations reported widely. Most likely this mutation could have been missed during most of the surveys due to selective genotyping/sequencing targeting three known *kdr* mutations. In most of the studies involving DNA sequencing, L1014 locus was not covered (Brownell et al., 2020; Durand et al., 2007; Firooziyan et al., 2017; Ghahvechi Khaligh et al., 2021; Komoda et al., 2020; Larkin et al., 2020; Ponce-Garcia et al., 2017; Roca-Acevedo et al., 2019; Toloza et al., 2014; Drali et al., 2012). It is therefore suggested that in all future studies, this mutation may be taken into account during sequencing.

This is the first study reporting any *kdr* mutation in Indian head louse populations and interestingly M827I showed the highest frequency of 19% and only 3% of T929I involved mainly in insecticide resistance. However, this study did not observe high frequency of these mutations as reported in studies from various countries (Hodgdon et al., 2010; Eremeeva et al., 2017; Mohammadi et al., 2021). Kerala population had the highest frequency of all the four mutations recorded in this study and only population with L1014F. There is urgent need for developing a cost-effective molecular method for genotyping all *kdr* mutations associated with pyrethroid resistance including L1014F, which will help in large scale surveillance of *kdr* mutations.

During this study, it was observed that the DNA sequencing of the VGSC encompassing IIS1-2 linker to IIS6 segments is challenging due to the frequent presence of indels in introns which lead to the collapse of sequence chromatogram (**Supplementary Figure S1**). Variation in the size of intron has also been reported by Lee et al. (2000). We, therefore, adapted a strategy to sequence VGSC using multiple primers designed from each exon. In cases where indels were present in the intron, it was not possible to get 2X sequence for all nucleotide positions.

In conclusion, this is the first study reporting the presence of all the three reported *kdr* mutations associated with pyrethroid resistance in head lice in Indian population along with the classical *kdr* mutation, L1014F, not reported earlier in this species.

## Authors’ contributions

Study design by OPS, experiments by PKM, AS, TG and SM, curation of data by AS, writing of the first draft by OPS and PKM. All authors read and approve the manuscript.

## Acknowledgments

We acknowledge the technical assistance rendered by Mr Uday Prakash.

## Research Integrity statement

The manuscript has been cleared by the Institutional Research Integrity Committee following ICMR Policy guidelines on Research Integrity and Publication Ethics (approval number 13/2022).

## Data availability

All data are incorporated in the article and its online supplementary material

## References

Amanzougaghene, N., Akiana, J., Mongo Ndombe, G., Davoust, B., Nsana, N. S., Parra, H. J., et al. (2016) Head Lice of Pygmies Reveal the Presence of Relapsing Fever Borreliae in the Republic of Congo. PLoS Neglected Tropical Disseases, 10, e0005142. doi: 10.1371/journal.pntd.0005142

Black WC, D. N. (1997) RAPD-PCR and SSCP analysis for insect population genetic studies In: The Molecular Biology of Insect Disease Vectors: a methods manual. Chapman & Hall, London.

Brownell, N., Sunantaraporn, S., Phadungsaksawasdi, K., Seatamanoch, N., Kongdachalert, S., Phumee, A., & Siriyasatien, P. (2020). Presence of the knockdown resistance (kdr) mutations in the head lice (Pediculus humanus capitis) collected from primary school children of Thailand. PLoS Neglected Tropical Diseases, 14(12), e0008955. doi: 10.1371/journal.pntd.0008955

Clark, J.M., Yoon, K.S., Lee, S.H. & Pittendrigh, B.R., (2013). Human lice: Past, present and future control. Pestic. Biochem. Physiol. 106, 162–171. doi: doi: 10.1016/j.pestbp.2013.03.008

Davies, T. G., Field, L. M., Usherwood, P. N., & Williamson, M. S. (2007). A comparative study of voltage-gated sodium channels in the Insecta: implications for pyrethroid resistance in Anopheline and other Neopteran species. Insect Molecular Biology, 16(3), 361–375. doi: 10.1111/j.1365-2583.2007.00733.x

Devore, C. D., Schutze, G. E., Council on School, H. & Committee on Infectious Diseases, A. A. o. P. (2015) Head lice. Pediatrics, 135, e1355–1365.

Dong, K., Du, Y., Rinkevich, F., Nomura, Y., Xu, P., Wang, L., Silver, K., & Zhorov, B. S. (2014). Molecular biology of insect sodium channels and pyrethroid resistance. Insect Biochemistry and Molecular Biology, 50, 1–17. doi: 10.1016/j.ibmb.2014.03.012

Drali, R., Benkouiten, S., Badiaga, S., Bitam, I., Rolain, J. M., & Brouqui, P. (2012). Detection of a knockdown resistance mutation associated with permethrin resistance in the body louse Pediculus humanus corporis by use of melting curve analysis genotyping. Journal of Clinical Microbiology, 50(7), 2229–2233. doi: 10.1128/JCM.00808-12

Durand, R., Millard, B., Bouges-Michel, C., Bruel, C., Bouvresse, S., & Izri, A. (2007). Detection of pyrethroid resistance gene in head lice in schoolchildren from Bobigny, France. Journal of Medical Entomology, 44(5), 796–798. doi: 10.1603/0022-2585(2007)44[796:doprgi]2.0.co;2

Eremeeva, M. E., Capps, D., Winful, E. B., Warang, S. S., Braswell, S. E., Tokarevich, N. K., et al. (2017) Molecular Markers of Pesticide Resistance and Pathogens in Human Head Lice (Phthiraptera: Pediculidae) From Rural Georgia, USA. Journal of Medical Entomology, 54, 1067–1072. doi: 10.1093/jme/tjx039

Firooziyan, S., Sadaghianifar, A., Taghilou, B., Galavani, H., Ghaffari, E. & Gholizadeh, S. (2017) Identification of Novel Voltage-Gated Sodium Channel Mutations in Human Head and Body Lice (Phthiraptera: Pediculidae). Journal of Medical Entomology, 54, 1337–1343. doi: 10.1093/jme/tjx107

Firooziyan, S., Sadaghianifar, A., Taghilou, B., Galavani, H., Ghaffari, E. & Gholizadeh, S. (2017) Identification of novel voltage-gated sodium channel mutations in human head and body lice (Phthiraptera: Pediculidae). Journal of Medical Entomology, 54, 1337–1343. doi: 10.1093/jme/tjx107

Ghahvechi Khaligh, F., Djadid, N. D., Farmani, M., Asadi Saatlou, Z., Frooziyan, S., Abedi Astaneh, et al. (2021). Molecular Monitoring of Knockdown Resistance in Head Louse (Phthiraptera: Pediculidae) Populations in Iran. Journal of Medical Entomology, 58(6), 2321–2329. doi: 10.1093/jme/tjab101

Ghahvechi Khaligh, F., Djadid, N. D., Farmani, M., Asadi Saatlou, Z., Frooziyan, S., Abedi Astaneh, et al. (2021). Molecular monitoring of knockdown resistance in head Louse (Phthiraptera: Pediculidae) Populations in Iran. Journal of Medical Entomology, 58(6), 2321–2329. doi: 10.1093/jme/tjab101

Hodgdon, H. E., Yoon, K. S., Previte, D. J., Kim, H. J., Aboelghar, G. E., Lee, S. H., et al. (2010) Determination of knockdown resistance allele frequencies in global human head louse populations using the serial invasive signal amplification reaction. Pest Managment Science, 66, 1031–1040. doi: 10.1002/ps.1979

Komoda, M., Yamaguchi, S., Takahashi, K., Yanase, K., Umezawa, M., Miyajima, A., Yoshimasu, T., Sato, E., Ozeki, R., & Ishii, N. (2020). Efficacy and safety of a combination regimen of phenothrin and ivermectin lotion in patients with head lice in Okinawa, Japan. The Journal of Dermatology, 47(7), 720–727. doi: 10.1111/1346-8138.15348

Larkin, K., Rodriguez, C. A., Jamani, S., Fronza, G., Roca-Acevedo, G., Sanchez, A., & Toloza, A. C. (2020). First evidence of the mutations associated with pyrethroid resistance in head lice (Phthiraptera: Pediculidae) from Honduras. Parasites & Vectors, 13(1), 312. doi: 10.1186/s13071-020-04183-2

Lee, S.H., Yoon, K.-S., Williamson, M.S., Goodson, S.J., Takano-Lee, M., Edman, J.D., Devonshire, A.L., Marshall Clark, J. (2000). Molecular analysis of kdr-like resistance in permethrin-resistant strains of head lice, Pediculus capitis. Pesticide Biochemistry & Physiology. 66, 130–143. doi: 10.1006/pest.1999.2460

Louni, M., Amanzougaghene, N., Mana, N., Fenollar, F., Raoult, D., Bitam, I., et al. (2018) Detection of bacterial pathogens in clade E head lice collected from Niger’s refugees in Algeria. Parasite & Vectors, 11, 348. doi: 10.1186/s13071-018-2930-5

Mohammadi, J., Azizi, K., Alipour, H., Kalantari, M., Bagheri, M., Shahriari-Namadi, M., et al. (2021) Frequency of pyrethroid resistance in human head louse treatment: systematic review and meta-analysis. Parasite, 28, 86. doi: 10.1051/parasite/2021083

Ponce-Garcia, G., Villanueva-Segura, K., Trujillo-Rodriguez, G., Rodriguez-Sanchez, I. P., Lopez-Monroy, B., & Flores, A. E. (2017). First detection of the kdr mutation T929I in head lice (Phthiraptera: Pediculidae) in School children of the Metropolitan Area of Nuevo Leon and Yucatan, Mexico. Journal of Medical Entomology, 54(4), 1025–1030. doi: 10.1093/jme/tjx045

Rinkevich, F. D., Du, Y. & Dong, K. (2013) Diversity and Convergence of Sodium Channel Mutations Involved in Resistance to Pyrethroids. Pesticide Biochemistry & Physiology, 106, 93–100. 10.1016/j.pestbp.2013.02.007

Robinson, D., Leo, N., Prociv, P., & Barker, S. C. (2003). Potential role of head lice, Pediculus humanus capitis, as vectors of Rickettsia prowazekii. Parasitology Research, 90(3), 209–211. doi: 10.1007/s00436-003-0842-5

Roca-Acevedo, G., Del Solar Kupfer, C. P., Dressel Roa, P., & Toloza, A. C. (2019). First determination of pyrethroid knockdown resistance alleles in human head lice (Phthiraptera: Pediculidae) From Chile. Journal of Medical Entomology, 56(6), 1698–1703. doi: 10.1093/jme/tjz101

Sasaki, T., Poudel, S. K., Isawa, H., Hayashi, T., Seki, N., Tomita, T., Sawabe, K., & Kobayashi, M. (2006). First molecular evidence of Bartonella quintana in Pediculus humanus capitis (Phthiraptera: Pediculidae), collected from Nepalese children. Journal of Medical Entomology, 43(1), 110–112. doi: 10.1093/jmedent/43.1.110

Sunantaraporn, S., Sanprasert, V., Pengsakul, T., Phumee, A., Boonserm, R., Tawatsin, A., et al. (2015) Molecular survey of the head louse Pediculus humanus capitis in Thailand and its potential role for transmitting Acinetobacter spp. Parasite & Vectors, 8, 127. 10.1186/s13071-015-0742-4.

SupYoon, K., Symington, S. B., Hyeock Lee, S., Soderlund, D. M. & Marshall Clark, J. (2008) Three mutations identified in the voltage-sensitive sodium channel alpha-subunit gene of permethrin-resistant human head lice reduce the permethrin sensitivity of house fly Vssc1 sodium channels expressed in Xenopus oocytes. Insect Biochemestry & Molecular Biology, 38, 296–306. doi: 10.1016/j.ibmb.2007.11.011

Toloza, A. C., Ascunce, M. S., Reed, D., & Picollo, M. I. (2014). Geographical distribution of pyrethroid resistance allele frequency in head lice (Phthiraptera: Pediculidae) from Argentina. Journal of Medical Entomology, 51(1), 139–144. doi: 10.1603/me13138

Tomita, T., Yaguchi, N., Mihara, M., Takahashi, M., Agui, N., & Kasai, S. (2003). Molecular analysis of a para sodium channel gene from pyrethroid-resistant head lice, Pediculus humanus capitis (Anoplura: Pediculidae). Journal of Medical Entomology, 40(4), 468–474. doi: 10.1603/0022-2585-40.4.468

Williamson, M. S., Martinez-Torres, D., Hick, C. A., & Devonshire, A. L. (1996). Identification of mutations in the housefly para-type sodium channel gene associated with knockdown resistance (kdr) to pyrethroid insecticides. Molecular & General Genetics: MGG, 252(1-2), 51–60. doi: 10.1007/BF02173204

