## Supplementary Figure S1 for "First report of classical knockdown resistance (*kdr*) mutation, L1014F, in human head louse *Pediculus humanus capitis* (Phthiraptera: Anoplura)"

**Supplementary materials**

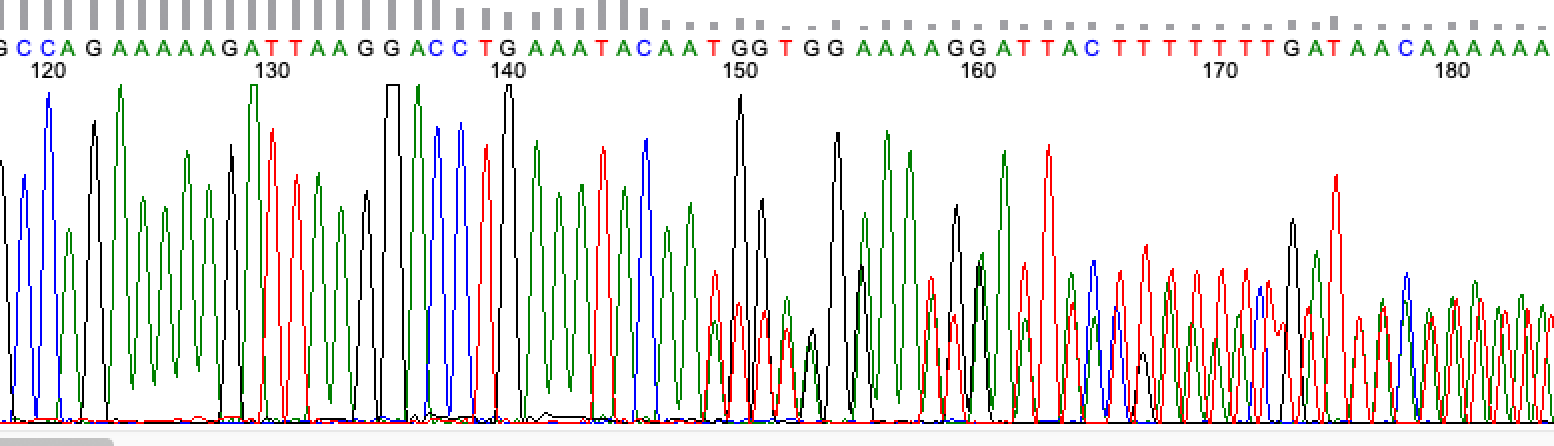

**Figure S1** A DNA sequence chromatogram (reverse direction) showing the collapse of sequence (from base position 149 onward) due to deletion of single ‘A’ nucleotide in the intron of one strand of DNA.
